# Selection on the FADS region in Europeans

**DOI:** 10.1101/086439

**Authors:** Matthew T. Buckley, Fernando Racimo, Morten E. Allentoft, Majken K. Jensen, Anna Jonsson, Hongyan Huang, Farhad Hormozdiari, Martin Sikora, Davide Marnetto, Eleazar Eskin, Marit E. Jørgensen, Niels Grarup, Oluf Pedersen, Torben Hansen, Peter Kraft, Eske Willerslev, Rasmus Nielsen

**Affiliations:** Departments of Integrative Biology and Statistics, University of California Berkeley, Berkeley, CA 94720, USA.; Natural History Museum of Denmark, University of Copenhagen, Øster Voldgade 5-7, 1350 Copenhagen K, Denmark.; The Novo Nordisk Foundation Center for Basic Metabolic Research, Section of Metabolic Genetics, Faculty of Health and Medical Sciences, University of Copenhagen, 2100 Copenhagen Ø, Denmark; National Institute of Public Health, University of Southern Denmark, 1353 Copenhagen, Denmark; Steno Diabetes Center, 2820 Gentofte, Denmark; Department of Computer Science, University of California, Los Angeles, CA 90095, USA; Department of Human Genetics, University of California, Los Angeles, CA 90095, USA; Program in Genetic Epidemiology and Statistical Genetics, Harvard T.H. Chan School of Public Health, Boston MA 02115, USA; Department of Nutrition, Harvard T.H. Chan School of Public Health & Channing Division of Network Medicine, Brigham and Women’s Hospital, Harvard Medical School, Boston MA 02115, USA; Department of Molecular Biotechnology and Health Sciences, University of Torino, Turin, Italy; Currently at Department of Genetics, Stanford University, Stanford, CA 94305, USA.

## Abstract

*FADS* genes encode fatty acid desaturases that are important for the conversion of short chain polyunsaturated fatty acids (PUFAs) to long chain fatty acids. Prior studies indicate that the *FADS* genes have been subjected to strong positive selection in Africa, South Asia, Greenland, and Europe. By comparing *FADS* sequencing data from present-day and Bronze Age (5-3k years ago) Europeans, we identify possible targets of selection in the European population, which suggest that selection has targeted different alleles in the *FADS* genes in Europe than it has in South Asia or Greenland. The alleles showing the strongest changes in allele frequency since the Bronze Age show associations with expression changes and multiple lipid-related phenotypes. Furthermore, the selected alleles are associated with a decrease in linoleic acid and an increase in arachidonic and eicosapentaenoic acids among Europeans; this is an opposite effect of that observed for selected alleles in Inuit from Greenland. We show that multiple SNPs in the region affect expression levels and PUFA synthesis. Additionally, we find evidence for a gene-environment interaction influencing low-density lipoprotein (LDL) levels between alleles affecting PUFA synthesis and PUFA dietary intake: carriers of the selected, derived allele have diminished increases in LDL cholesterol with a higher intake of PUFAs. We hypothesize that the selective patterns observed in Europeans were driven by a change in dietary composition of fatty acids following the transition to agriculture, resulting in a lower intake of arachidonic acid and eicosapentaenoic acid, but a higher intake of linoleic acid and α-linolenic acid.

## Introduction

Long-chain polyunsaturated fatty acids (LC-PUFAs) are important components of mammalian tissue and are crucial for a variety of biological processes. They are bioactive elements of cell membranes and have an important role in neuronal membrane development (Marszalek and Lodish 2005; Darios and Davletov 2006). With large brains composed mostly of lipids, humans have a particularly strong requirement for these fatty acids (Mathias, *et al*. 2012). LC-PUFAs also serve as precursors for cell signaling molecules including eicosanoids, such as prostaglandins, which act as messengers in the central nervous system and exert control over many bodily systems including inflammation (Hester, *et al*. 2014). LC-PUFA concentration has been linked to infant visual and brain development (McCann and Ames 2005) and to risks of cardiovascular and coronary heart disease and mortality (Patel, *et al*. 2010). The physiologically most important LC-PUFAs include ω-6 (or n-6) PUFA arachidonic acid (ARA; 20:4n-6), and ω-3 (or n-3) PUFAs eicosapentaenoic acid (EPA; 20:5n-3) and docosahexaenoic acid (DHA; 22:6n-3). Additionally, ω-3 LC-PUFAs are common dietary supplements, despite ongoing debate regarding their potential preventative role against cancer or heart disease (MacLean, *et al*. 2006; Rizos, *et al*. 2012).

Humans can obtain LC-PUFAs directly, particularly by the consumption of meat, fish, and marine mammals. Alternatively, these compounds can be synthesized endogenously from the short chain ω-6 and ω-3 polyunsaturated fatty acids (SC-PUFAs) linoleic acid (LA; 18:2n-6) and α-linolenic acid (ALA; 18:3n-3), (Figure 1). These SC-PUFAs are considered essential and are obtained primarily through the consumption of vegetable oils. The rate-limiting steps in the synthesis of long chain PUFAs from short chain PUFAs are catalyzed by two fatty acid desaturases: delta-5 desaturase (D5D) and delta-6 desaturase (D6D) (Cho, *et al*. 1999), encoded by *fatty acid destaurase 1* (*FADS1*) and *fatty acid desaturase 2* (*FADS2*), respectively. *FADS1* and *FADS2* are located adjacent to each other on chromosome 11 (11q12-13.1), and next to another gene encoding a third fatty acid desaturase (*FADS3*). The function of *FADS3* has yet to be elucidated, but the gene shares 52% and 62% sequence homology with *FADS1* and *FADS2*, respectively, and likely resulted from gene duplication (Marquardt, *et al*. 2000).

**Figure 1.**
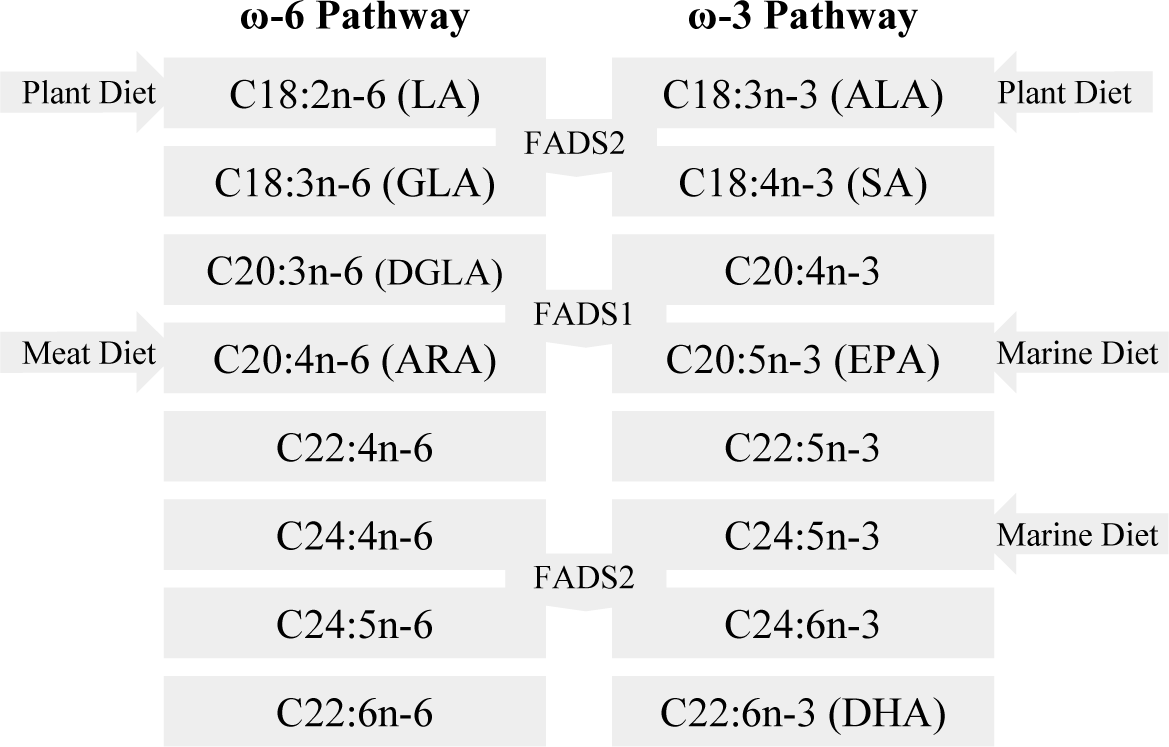
n-3 and n-6 polyunsaturated fatty acid synthesis pathway. The main dietary intakes are shown and fatty acid desaturases (FADS1 and FADS2) are shown in context of the fatty acids they directly affect.

Previous studies have suggested that the *FADS* genes are targets of natural selection in different human populations. Mathias *et al*. (2011; 2012) compared allele frequencies of 80 *FADS* SNPs across African and non-African populations, showing that African populations had substantially higher frequencies of derived alleles associated with more efficient desaturases (Mathias, *et al*. 2011). In a subsequent study, Ameur *et al*. (2012) showed that the *FADS* genes span two distinct linkage disequilibrium (LD) blocks, one of which contains an ancestral haplotype largely present in Eurasia, and a derived haplotype that is prevalent in Africa. Both these studies argued that a selective sweep began in the *FADS* genes prior to the human migration out of Africa, roughly 85 thousand years ago (kya). Humans migrating out of Africa putatively carried mostly the ancestral haplotype, which remained in high frequency in non-African populations, while the derived haplotype came close to fixation in Africa. It is unclear why positive selection for the derived haplotype appears to be restricted to Africa. Mathias *et al*. (2012) suggested that the emergence of regular hunting of large animals, dated to approximately 50 thousand years ago (kya), might have diminished the pressure for humans to endogenously synthesize LC-PUFAs.

In a subsequent genome-wide scan for positive selection in the Greenlandic Inuit, the *FADS* region was found to be the strongest outlier region based on patterns of allele frequency differentiation relative to other populations (Fumagalli, *et al*. 2015). The two most highly differentiated SNPs (rs7115739 and rs174570), both located in *FADS2*, were associated with decreased concentrations of LC-PUFAs but increased concentrations of SC-PUFAs (Fumagalli, *et al*. 2015). In their traditional diet, Inuit consume extremely high levels of LC-PUFAs from fish and marine mammals, ostensibly diminishing their need to endogenously synthesize these LC-PUFAs, but have a low intake of some SC-PUFAs such as linoleic acid. The mutations in the Inuit appear to compensate for a decreased intake of SC-PUFAs and an increased intake of LC-PUFAs. Fumagalli *et al*. (2015) also demonstrated that the selected alleles were associated with a decrease in weight, height, fasting serum insulin, and fasting serum LDL cholesterol. The *FADS* genes have been associated with metabolic traits in multiple previous studies (Bokor, *et al*. 2010; Glaser, *et al*. 2011), but the strong association with height had not been noted previously, presumable because these SNPs segregate at very low frequencies in Europeans. Nevertheless, the effect on height was replicated in European cohorts (Fumagalli, *et al*. 2015).

More recently, Kothapalli *et al*. (2016) studied genomes from populations in South Asia and showed strong signs of positive selection for an indel (rs66698963, mislabeled as rs373263659 in 1000 Genomes Project Phase III data) in *FADS2*, with the insertion allele putatively endowing South Asians with the ability to more efficiently synthesize LC-PUFAs, possibly as an adaptation to a more vegetarian diet.

Mathieson *et al*. (2015) carried out a genome-wide scan for positive selection comparing DNA microarray data from ancient and present-day European genomes. They carried out the selection scan using a linear model aimed at predicting present-day allele frequencies from the allele frequencies in ancient source populations from the European Neolithic and Bronze Ages. They then identified SNPs with allele frequencies that strongly deviated from those predicted by the genome-wide pattern. One of these SNPs was rs174546, the derived allele (C) of which tags the derived haplotype in Africans (Ameur, *et al*. 2012) and is in high LD (r^2^ = 0.978; CEU) with the derived allele of rs174537, a SNP located in the middle of the selection peak in Mathias *et al*. (2012). The Mathieson *et al*. (2015) study provides strong evidence of selection in the *FADS* region in Europe over the past 4,000 years, in addition to the patterns of selection already reported in Africans, South Asians, and the Inuit.

The aim of this study is to further investigate the selection signal in Europeans and the functional effects of the selected alleles. Taking advantage of genome-wide sequencing data from 101 Bronze Age individuals by Allentoft *et al*. (2015) as well as data from the 1000 Genomes Project (Genomes Project, *et al*. 2015), we identify novel potential targets of selection in the *FADS* region. Using expression data and data from multiple GWASs, we investigate the functional effects of the allelic variants that have increased in frequency in Europe over the past 5,000 years. We show that SNPs associated with increased expression of *FADS1* and increased production of arachidonic acid and eicosapentaenoic acid have been favored in Europeans since the Bronze Age. Our results suggest that selection in the *FADS* region is complex and has targeted several loci across different populations.

## Results

### Origin and structure of human haplotypes in the FADS region

As previously observed (Ameur, *et al*. 2012; Mathieson, blog post) there are two distinct LD blocks spanning the FADS gene cluster. These blocks are especially evident when focusing on Europeans (Figure. 3, Suppl. Fig. 1) The first block (Block 1: chr11:61547000-61625000) overlaps *FADS1* and half of *FADS2*, while the second block (Block 2: chr11:61627000-61673000) overlaps with the rest of *FADS2* and all of *FADS3* (Figure 2).

**Figure 2.**
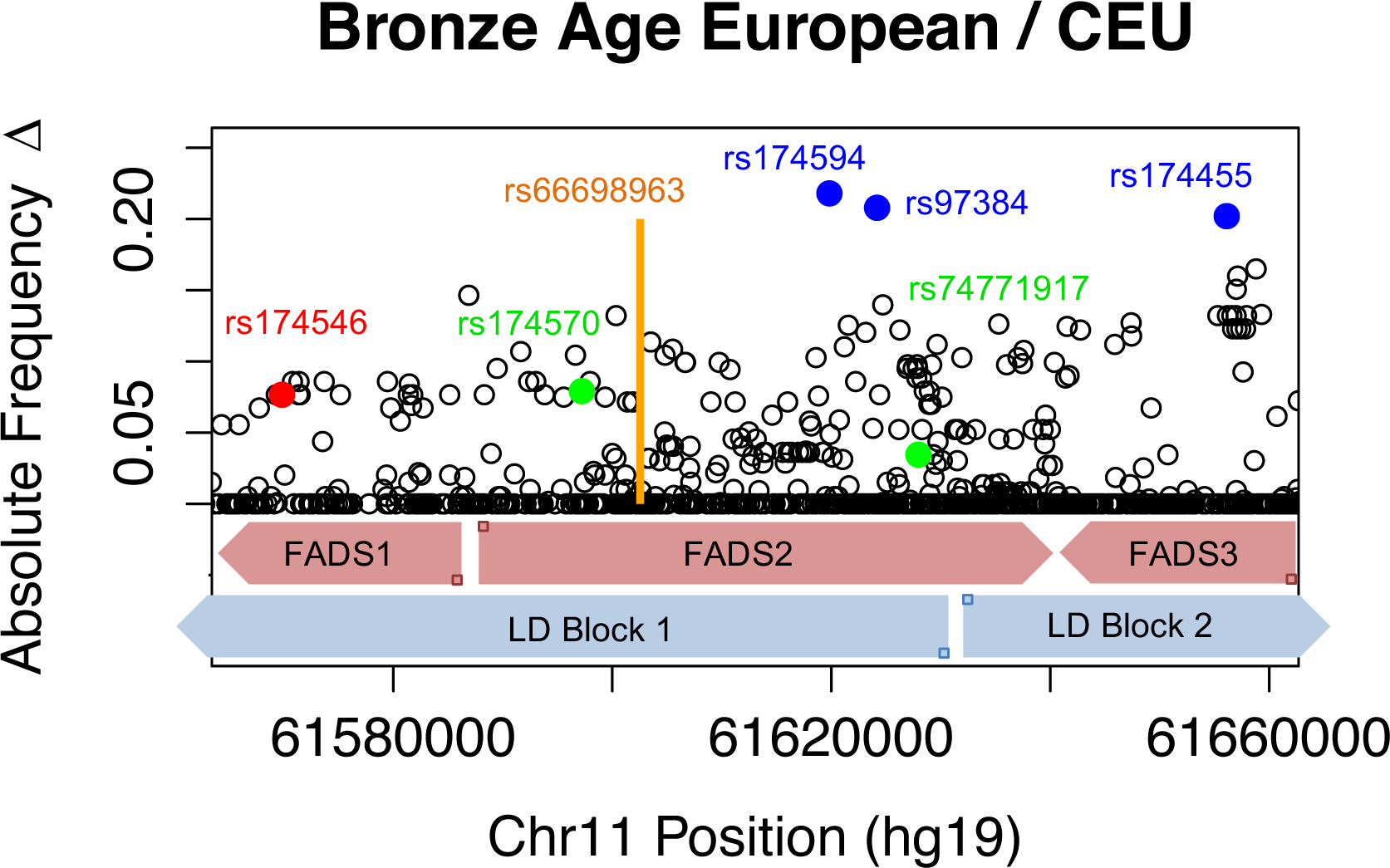
Allele Frequency Changes across *FADS* region. Four SNPs exhibiting the greatest change are labeled as well as rs174546, which has previously shown signs of positive selection. The approximate location of *FADS1* is in blue, *FADS2* in purple, *FADS3* in green. The x-axis depicts the position on chromosome 11 using coordinates from build Hg19.

We set out to analyze the selective processes in Europe in detail, across both space and time. First, we inspected the haplotype structure at this locus using present-day human genomes from phase 3 of the 1000 Genomes Project (Genomes Project, *et al*. 2015). As previously noted, the *FADS* cluster shows high amounts of haplotypic variation in present-day humans (Mathias, *et al*. 2012; Mathieson, blog post). In Block 1, this variation is largely attributable to high differentiation between two haplotype clusters: a cluster widespread in Africa, largely containing derived alleles and possibly subject to a selective sweep (Mathias, *et al*. 2011; 2012), and an ancestral cluster, which is present across Eurasia. Interestingly, two archaic human genomes (Altai Neanderthal and Denisova) that are sister groups to each other genome-wide, actually cluster with different clusters in this region (Figure 3, Suppl. Figures 2, 3): the Denisovan genome clusters with the ancestral (Eurasian-specific) cluster, while the Altai Neanderthal genome clusters with the derived cluster, which is prevalent in Africa. Mathieson (blog post 2015) argues this pattern could be explained by a re-introduction of the ancestral cluster into Eurasians via introgression from archaic humans, followed by a second selective event in Eurasia.

It is worth noting, however, that this locus does not show signatures that could be consistent with a simple model of introgression, at least directly from the populations to which the sequenced archaic genomes belong. Unlike other documented cases of adaptive introgression (Racimo, Gokhman, et al. 2016) inspecting of the haplotype network reveals that none of the branches connecting the archaic haplotypes to their most similar present-day human haplotypes are less than half as large as the branches connecting those same present-day haplotypes to other present-day human haplotypes (Figure 3). Additionally, we observe no archaic alleles in this region that are at low frequency (<1%) in Africans but at high (>20%) frequency in any particular non-African population or continental panel from the 1000 Genomes data – a signature of archaic adaptive introgression (Racimo, Marnetto, et al. 2016). Finally, this region does not appear as a significant candidate in recent scans for archaic adaptive introgression in Eurasians (Sankararaman, *et al*. 2014; Vernot and Akey 2014; Sankararaman, *et al*. 2016). It is, therefore, possible that the two haplotype clusters perhaps reflect ancient balancing selection rather than introgression, although we cannot fully discard complex scenarios with multiple introgression and/or selective sweep events in both Africa and Eurasia.

**Figure 3.**
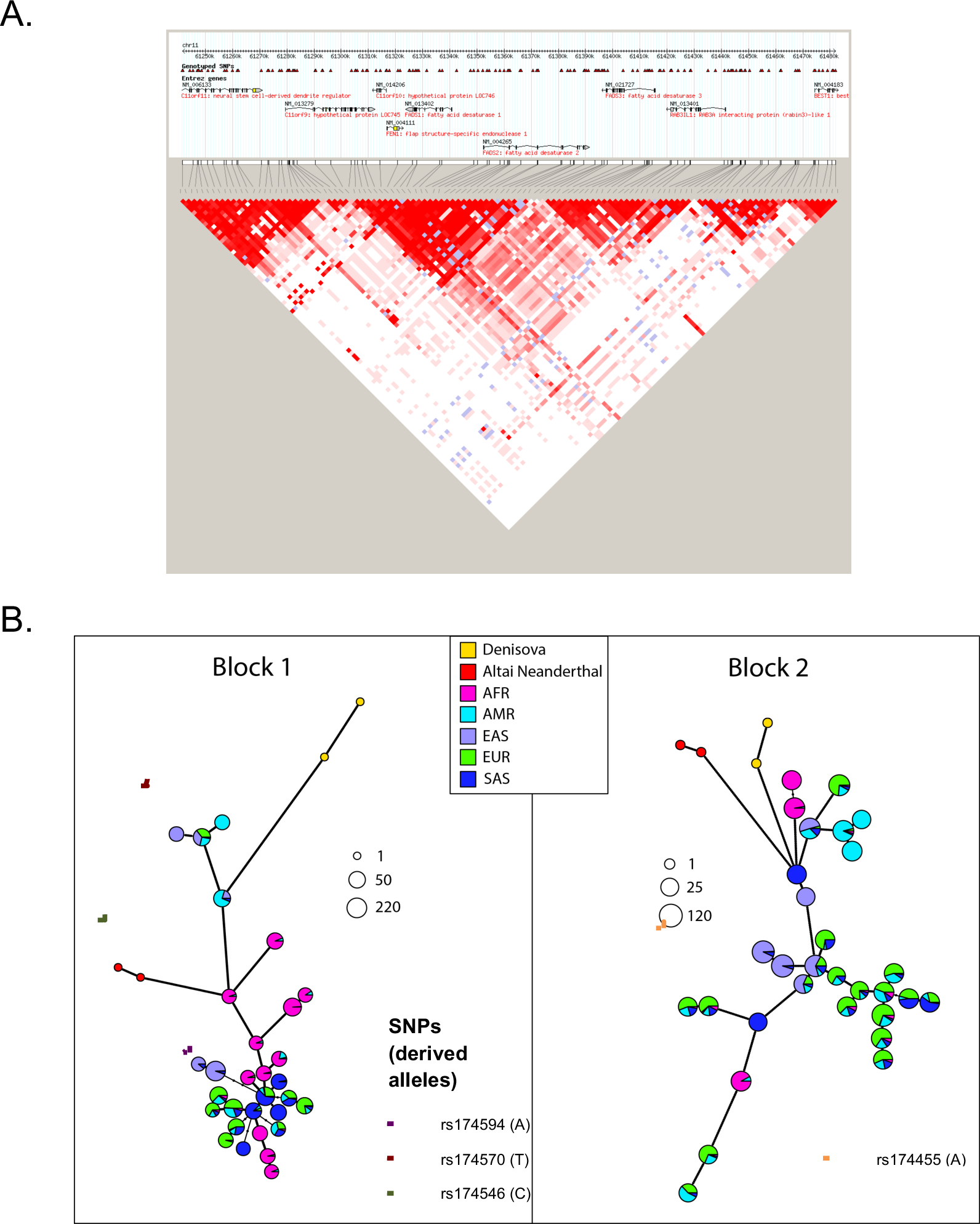
A) Linkage disequilibrium plots for the FADS region for CEU+TSI. Each colored cell corresponds to a SNP pair. For SNP pairs with LOD scores smaller than 2, values of D’ = 1 are denoted in blue, while values of D’ < 1 are denoted in white. For SNP pairs with LOD scores larger than 2, values of D’ are represented in a pink-to-red color scheme, with stronger shades of red corresponding to higher values of D’. Note that the SNP positions in the top scale of each LD plot are in hg18 coordinates. B) Haplotype network plots of the two LD blocks in the *FADS* region, using the phase 3 1000 Genomes data, built using pegas (Paradis *et al*. 2010). Block 1: chr11:61547000-61625000. Block 2: chr11:61627000-61673000. The colors denote the continental populations to which each haplotype belongs. The size of the pie charts is proportional to log2(*n*) where *n* is the number of individuals carrying the haplotype. The black dots on each connecting line denote the number of differences separating each haplotype from its neighbors. The dotted lines denote the haplotype clusters in which each of the putatively derived alleles from four interesting SNPs are located: rs174570 (Fumagalli *et al*. 2015), rs174546 (from Mathieson’s blog post), rs174594 (HDS from this study) and rs174455 (HDS from this study).

We also aimed to determine where in the haplotype network each of the derived alleles from the SNPs discussed in this study were located (Figure 3, Suppl. Fig. 3). We find that the derived allele (T; 16%, CEU) of rs174570 – which showed the strongest signs of selection in the Greenlandic Inuit – is present in a non-African haplotype cluster in Block 1, but not present in Africans or the archaic humans. The derived allele (C; 64%, CEU) from the putatively selected SNP in Mathieson’s blog post, rs174546, is in the other haplotype cluster, which includes several African and non-African haplotypes, as well as the Altai Neanderthal genome. The Denisovan genome and the other non-African haplotypes carry the ancestral allele at this site (Figure 3, Suppl. Fig. 2). The derived allele of rs174570 is present in a different cluster of non-African haplotypes in Block 1, which are closer to African haplotypes. Finally, the derived allele of rs174455 (A; 65%, CEU) is present in a mostly non-African haplotype cluster in Block 2.

### Allele frequency changes in Europe

We then set out to compare patterns of temporal allele frequency differentiation in Europe, by comparing the allele frequencies of *FADS* SNPs in present-day CEU from the 1000 Genomes Project (Genomes Project, *et al*. 2015) and 56 Bronze Age Europeans (Allentoft, *et al*. 2015). The vertical axis of Figure 2 shows the absolute value of allele frequency changes across the *FADS* cluster. Below, we refer to the four top SNPs (rs174594, rs97384, 174455 and rs174465. See Table 1.), each with >17% change in allele frequency, as highly differentiated SNPs (HDSs) and they are labeled in blue in Figure 2. rs174594 and rs97384 are located in *FADS2*, towards the end of LD Block 1, and both are in high LD (*r*^2^ = 0.917) in the 1000 Genomes CEU panel. rs174455 and rs174465 are located 2.7 kb apart in *FADS3* on LD Block 2, and are also in relatively high LD (*r*^2^ = 0.776, CEU) with each other, but less so with rs174594 and rs97384 (*r*^2^ ≈ 0.4). The lead SNPs from our study were not included in the SNP typing platform used by Mathieson, *et al*. (2015) and may therefore have been missed in that study.

**Table 1.** SNP selected allele frequencies reference

| SNP | Ancestral Allele | Derived Allele | Allele under putative positive selection in Europe | Frequency of Selected Allele in Bronze Age Europeans | Frequency of Selected Allele in 1KGP CEU |
| --- | --- | --- | --- | --- | --- |
| rs174546 | T | C | C | 51% | 64% |
| rs66698963 | - | + | - | No Data | 47% |
| rs174594 | C | A | A | 39% | 62% |
| rs97384 | T | C | C | 40% | 61% |
| rs174455 | G | A | A | 45% | 65% |
| rs174465 | C | T | T | 52% | 70% |

In addition to CEU, we also compared two other groups comprised of panels from the 1000 Genomes Project (Genomes Project, *et al*. 2015), Northern European (FIN, GBR, CEU) and Southern European (TSI, IBS), to the Bronze Age data from Allentoft, *et al*. (2015) (Figure 4). The ancestral haplotype in LD Block 1 appears to be more prevalent in Northern Europeans and Bronze Age Europeans than in Southern Europeans, resulting in lower allele frequency changes in the comparison between Northern and Bronze Age Europeans (Figure 4.A) than in the comparison between Southern and Bronze Age Europeans (Figure 4.B). The lead SNP from Mathieson, *et al*. (2015) (rs174546) in LD Block 1 is one of most highly differentiated SNPs between present-day Northern and Southern Europeans (Figure 4.C), whereas our HDSs show less differentiation between the present-day populations. This suggests LD Block 1 may have been subject to differential selection pressures in different European populations at different points in time and this may drive some of the signal observed in Mathieson, *et al*. (2015). Alternatively, the allele frequency difference between Northern and Southern Europe could be driven by differential contributions of Neolithic DNA originating from the Middle East in these two groups.

**Figure 4.**
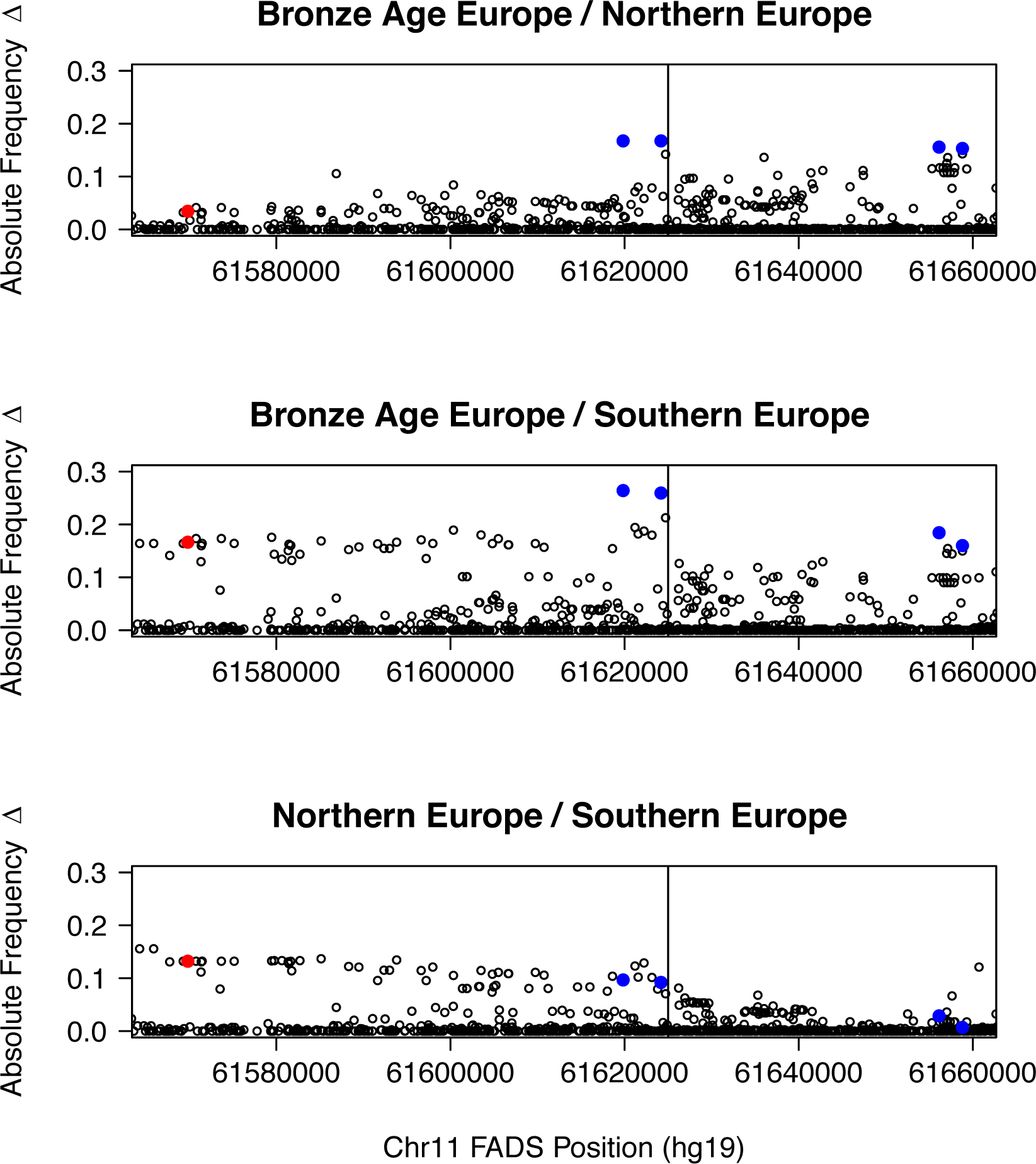
Comparisons of Bronze Age Europe to present-day Northern Europe (1000 Genomes Project CEU, FIN, GBR) and Southern Europe (IBS, TSI). Note the string of SNPs in Southern Europeans that show greater allele frequency change. The vertical line in both plots represents the approximate transition point between LD Block 1 and LD Block 2.

### Comparison of targets of selection in South Asia, Europe, and Greenland

In Figure 2, we also show lead SNPs and indels from other previous population genetic studies of the *FADS* region: rs174546 from Alexander, *et al*. (2009); Mathieson, *et al*. (2015) (red), two SNPs from Fumagalli, *et al*. (2015) (green), and rs66698963 from Kothapalli, *et al*. (2016) (orange). Note that the Allentoft *et al*. (2015) low-coverage aDNA dataset does not include called indels so we did not plot the allele frequency change for rs66698963. rs174546, which also tags the Ameur, *et al*. (2012) derived haplotype in Africans is additionally in very high LD (r^2^=0.978, CEU) with the lead SNP from Mathias, *et al*. (2012) (rs174537). These two SNPs and the two HDSs on LD Block 1 (rs174594, rs97384) are tightly linked to each other in Europe (Suppl. Table 1). The indel found to be under selection in South Asians (rs66698963) (Kothapalli, *et al*. 2016), the top hit in the Greenlander Inuit scan (rs174570) (Fumagalli, *et al*. 2015), and HDS rs174455 are not in exceptionally high LD with each other or with rs174546 (Suppl. Table 1)

We also noticed substantial differences in LD patterns among populations, as can be observed in Supplementary Table 1. For example, rs66698963 is in relatively strong LD with all other SNPs in the Bengali from Bangladesh (BEB), but much less so in other populations, presumably due to the selective pressures acting in South Asia and described by Kothapalli, *et al*. (2016) (Suppl. Fig. 1). In contrast, it is in much lower LD with the other SNPs in CEU: the *r*^2^ value in CEU between the lead SNP of this study (rs174594) and rs66698963 is only 0.12. rs66698963 is, therefore, unlikely to be the target of selection in Europe. Similarly, the *r*^2^ value between rs174570, the lead SNP of the Inuit study (Fumagalli, *et al*. 2015), and rs174594 and rs66698963 is only 0.31 and 0.06, respectively, suggesting that the selective pressures acting on the *FADS* region have had different targets in Europeans, South Asians and the Inuit.

### Identifying targets of selection in Europe

To further determine whether one of the lead SNPs from this study or rs174546 is the most likely target of selection in Europe, we compared allele frequencies before and after selection using a likelihood approach (Supplementary Information 2), jointly modeling selection affecting one or more SNPs. We use rs174594 as the lead SNP to represent the SNPs from the present study.

The results of the analyses are shown in Supplementary Information 2, Tables 1 and 2. We compared three models: a model in which selection acts only on the derived allele (C) of rs174546 (Model 1), a model in which it only acts on the derived allele (A) of rs174594 (Model 2), and a model in which a haplotype (DD) carrying the derived alleles from both rs174546 and rs174594 has been the target (Model 3). Both model 2 and 3 have likelihood values that are approx. 10 likelihood units better than Model 1. This suggests that the change in allele frequency is best modeled as driven either by selection targeting the DD haplotype or by selection favoring the derived allele in rs174594 alone. Selection favoring only the derived allele in rs174546 (Model 1) is not a parsimonious explanation of the data. However, we cannot confidently distinguish between selection favoring the DD haplotype versus selection acting only on rs174594, although a model in which selection acted on rs174594 has a slightly higher likelihood than the model of selection acting on DD.

### Expression Associations

To investigate possible functional effects of the selected haplotypes/alleles in Europe, we examined public data from the Genotype-Tissue Expression (GTEx) project (Lonsdale, *et al*. 2013) for the four HDSs as well as rs174546 (Table 2; Suppl. Tables 3A,3B) The most significant results are in whole blood, where all SNPs are eQTLs for *FADS2* with p-values < 10^-19^. In all cases, the effect size (linear regression coefficient, units of quantile normalized gene expression over allele dosage) of the derived allele (corresponding to the allele that has increased frequency over time) is to decrease *FADS2* expression. The largest effect sizes and smallest p-values are for rs174546.

**Table 2.** Abbreviated GTEx eQTL Results, p-value cut off of 10^-10^

**(A) Effect on FADS2 Expression**
| SNP | p-value | Derived Allele Effect Size | Tissue |
| --- | --- | --- | --- |
| rs174546 | 4.00E-29 | -0.77 | Whole Blood |
| rs174594 | 2.90E-21 | -0.63 | Whole Blood |
| rs97384 | 9.30E-20 | -0.62 | Whole Blood |
| rs174546 | 5.80E-13 | -0.26 | Cells - Transformed fibroblasts |
| rs174546 | 8.80E-12 | -0.42 | Thyroid |
| rs174465 | 3.80E-11 | -0.46 | Whole Blood |
| rs174546 | 3.90E-11 | -0.5 | Esophagus - Muscularis |
| rs174455 | 6.20E-11 | -0.45 | Whole Blood |
| rs174594 | 9.40E-11 | -0.46 | Esophagus - Muscularis |
| rs174546 | 1.00E-10 | -0.37 | Colon - Transverse |
| rs97384 | 3.10E-10 | -0.46 | Esophagus - Muscularis |
| rs174594 | 5.50E-10 | -0.21 | Cells - Transformed fibroblasts |

**(B) Effect on FADS1 Expression**
| SNP | p-value | Derived Allele<br>Effect Size | Tissue |
| --- | --- | --- | --- |
| rs174546 | 1.40E-15 | 0.73 | Pancreas |
| rs174546 | 2.30E-12 | 0.72 | Brain - Cerebellum |
| rs174594 | 1.10E-10 | 0.6 | Pancreas |
| rs174546 | 4.60E-10 | 0.41 | Esophagus - Mucosa |
| rs97384 | 5.90E-10 | 0.6 | Pancreas |

There are also expression associations for *FADS1*, with the smallest p-values found in the pancreas (p < 10^-9^ for rs174546, rs174594, rs97384). Interestingly, the effects sizes are in the opposite direction as those for *FADS2*; the derived alleles are associated with an increase in expression for *FADS1*. This associated increase in FADS1 expression is also consistent with the results by Wang, et al. (2015) across 154 human liver tissue samples. Although the effect of these expression changes for LC-PUFA synthesis is somewhat difficult to predict without further data, we expect this would lead to a likely increase in arachidonic acid and eicosapentaenoic acid concentrations. We also note that *FADS1* and *FADS2* are transcribed in opposite directions and share their promoter regions. It is possible that the opposite directed effects on expression are caused by interference in the promoter regions of the two genes due to structural requirements or competition of transcription factors.

### Allelic Heterogeneity

To examine if the expression data also show evidence for an effect of multiple alleles, mirroring that observed for allele frequency changes, we also applied an extension of CAVIAR (Hormozdiari, *et al*. 2014), a probabilistic method developed to determine a set of all causal variants in a given locus while allowing that more than one variant may be causal. The extended version of CAVIAR thus allows for the detection of allelic heterogeneity (AH). We observed AH with high confidence for *FADS1* expression in tibial artery tissue (Pr(2 causal variants) = 0.65, full results in Supplementary Table 2). For *FADS2* expression, we observed evidence of AH in three tissue types: transformed fibroblast cells (Pr(2 causal variants) = 0.72), left heart ventricle (Pr(2 causal variants) = 0.74), and whole blood (Pr(3 causal variants) = 0.74). Tests for *FADS3* did not reveal strong evidence of AH. The inferred AH associated with *FADS2* in whole blood observed here, taken together with the previously described results on allele frequency changes, suggest that there are multiple targets of selection in the FADS genes.

### Functional characterization of HDSs

The strong expression associations may suggest that one or more of the investigated variants have a regulatory function, in addition to the previously proposed function of the rs66698963 indel (Reardon, *et al*. 2012; Kothapalli, *et al*. 2016). Therefore, we investigated the genomic features of the HDSs using CADD (Kircher, *et al*. 2014) and the WashU Epigenomics Browser (Zhou, *et al*. 2015). All of the SNPs are in either intronic or intergenic regions. rs174546 is located in an intron of *FADS1* and has a high GERP score (4.54) (Davydov, *et al*. 2010) suggesting strong past purifying selection in this site. rs174595 is located in an intron of *FADS2* and is also in a Segway segment (Hoffman, *et al*. 2012) associated with transcription factor activity. rs174455 flanks an active transcription start site region of *FADS3*. Finally, rs174465 is in the promoter of *FADS3*. Though some of these sites are located in interesting genomic features, it still remains unclear which are causal. All annotations are listed in Suppl. Table 4.

### Phenotypic associations in Global Lipids Genetics Consortium GWAS

There are multiple known associations between SNPs in the *FADS* region and lipid- and metabolism-related phenotypes (Mathias, *et al*. 2011; Ameur, *et al*. 2012; Hester, *et al*. 2014; Fumagalli, *et al*. 2015). The Global Lipids Genetics Consortium (2013) (GLGC) carried out the most comprehensive GWAS study for loci that influence lipid metabolism, using genomic data from 188,577 individuals of primarily European ancestry. The GWAS study specifically examined high-density lipoprotein cholesterol (HDL), low-density lipoprotein cholesterol (LDL), total cholesterol (TC), and triglyceride (TG) phenotypes. Only 4 loci, out of 157 loci associated with lipid levels at P ≤ 5 x 10^-8^, were associated with effects on all 4 phenotypes (HDL, LDL, TC, and TG). One of these 4 loci was the *FADS* gene cluster. We used LocusZoom (Pruim, *et al*. 2010) to generate plots with the publically available GLGC GWAS data.

Figure 5 includes 3 such plots depicting GWAS results for LDL while plots for HDL, TC and TG can be found in Suppl. Fig. 4. In Figure 5, panel A is centered on rs174546 in *FADS1*, while panel B and panel C are centered around an HDS from the remaining genes, rs97384 in *FADS2* and rs174455 in *FADS3*. The panels show approximate LD relationships by color-coding the markers’ r^2^ values with respect to the labeled SNP in the center of each plot. Of these three SNPs, rs174546 shows the strongest phenotypic associations, specifically linking the presence of the derived allele (C) with increased levels of LDL, HDL, TC and decreased levels of TG (p-values 2x10^-39^, 8x10^-28^, 3x10^-37^, 7x10^-38^). rs174546 also showed the greatest LD with neighboring SNPs, despite the fact that rs97384 and rs174455 have undergone stronger changes in allele frequencies over the last 4,000 years.

**Figure 5.**
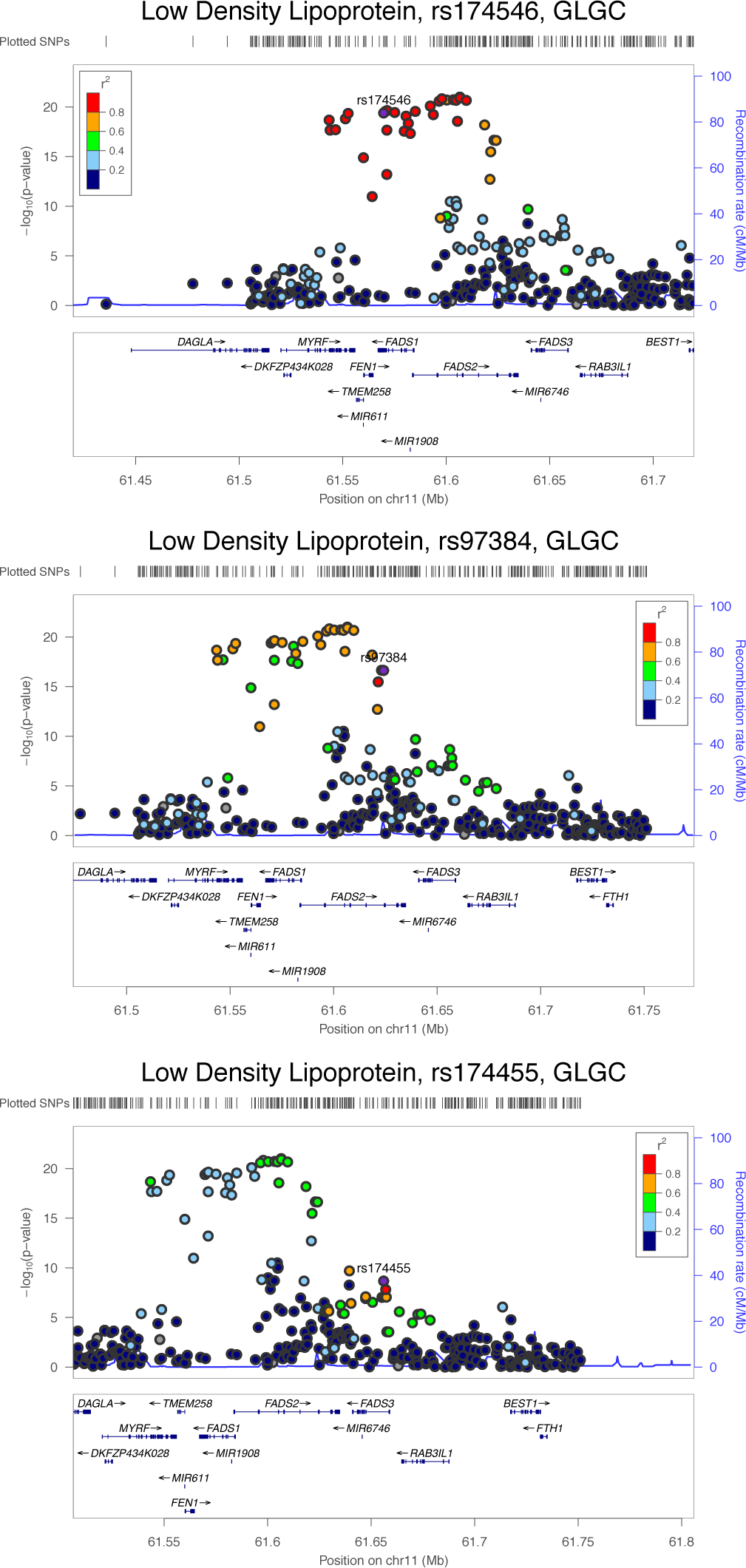
LocusZoom plots for 3 SNPs from the Global Lipids Genetics Consortium joint GWAS for high-density lipoprotein (HDL) cholesterol levels. We note in (A) that rs174546 in *FADS1* is the most significant SNP across the entire region. It is also in high LD with all other similarly significant SNPs in the region. (B) includes SNP rs97384 from *FADS2*, which despite its relatively large allele frequency change, is not particularly significant in the GWAS. (C) shows a plot for rs174455, the most prominent SNP in *FADS3*.

### Phenotypic associations in ADDITION-PRO Metabolomics Study

To further investigate phenotypic associations, we queried the Danish ADDITION-PRO study (Johansen, et al. 2012), which contains data from 2082 participants for 231 lipid-related phenotypes obtained at 3 time points (after fasting and 30 minutes and 120 minutes after oral intake of 75g glucose) as part of an oral glucose tolerance test (Table 3; Suppl. Table 5). The database does not contain LC-PUFAs, but it does contain a phenotype related to SC-PUFAs: linoleic acid to total fatty acids ratio, and this phenotype is in fact the phenotype for which we find the strongest association. For both rs174594 and rs174546 the derived alleles are associated with a decrease in the level of linoleic acid (p < 10^-12^, Bonferroni-corrected significance threshold 2.405x10^-5^). The second most significant phenotypic association (p < 10^-11^) was between rs174546 and the “estimated degree of unsaturation.” In this case, the effect of the derived allele was to increase the degree of unsaturation, which is consistent with the notion that the derived allele promotes rapid conversion of SC-PUFAs (such as linoleic acid) to LC-PUFAs.

**Table 3.** Top Associations from AdditionPRO Metabolomics GWAS

| SNP | P-Value | Derived Allele<br>Effect Size | Phenotype |
| --- | --- | --- | --- |
| rs174546 | 2.71E-13 | -0.924 | Linoleic acid / Fatty Acids (Fasting) |
| rs174546 | 2.98E-13 | -0.9114 | Linoleic acid / Fatty Acids (30 min) |
| rs174594 | 9.16E-13 | -0.9418 | Linoleic acid / Fatty Acids (Fasting) |
| rs174546 | 2.7E-12 | 0.02137 | Estimated degree of unsaturation (120 min) |
| rs174546 | 4.44E-12 | -0.8802 | Linoleic acid / Fatty Acids (120 min) |
| rs174594 | 4.72E-12 | -0.902 | Linoleic acid / Fatty Acids (30 min) |
| rs174594 | 2.43E-11 | -0.8874 | Linoleic acid / Fatty Acids (120 min) |
| rs174594 | 8.77E-10 | 0.01948 | Estimated degree of unsaturation (120 min) |
| rs174546 | 9.7E-10 | 0.01726 | Estimated degree of unsaturation (Fasting) |
| rs174546 | 9.85E-10 | 0.01749 | Estimated degree of unsaturation (30 min) |

### Phenotypic associations and gene-diet interactions in Nurses’ Health Study and the Health Professionals Follow-up Study

An analysis of the combined Nurses’ Health Study (NHS) and Health Professionals Follow-Up Study (HPFS) GWAS results yielded further evidence of phenotypic associations. The analysis of biological samples included measurements of LC-PUFAs (EPA and ARA), total cholesterol and lipoprotein concentrations, as well as key metabolites and products directly influenced by *FADS* activity. We queried our 4 HDSs and found the strongest associations with rs174594 for different fatty acid phenotypes (see Suppl. Table 6). The derived allele of rs174594 showed positive associations with plasma EPA levels (beta = 0.096 increase in concentration [measured as percent of total fatty acids] per derived allele [95% CI 0.069-0.12], p value 1.8x10^-11^), plasma ARA levels (beta = 1.18 increase in concentration per derived allele [95% CI 1.06-1.30], p value 3.6x10^-102^), as well as higher plasma total and LDL cholesterol levels (beta = 3.50 mg/dL per derived allele [95% CI 0.99-6.01] for total cholesterol and 3.03 mg/dL [95% CI 0.76-5.30], p value = 0.006 and 0.009, respectively). These results reinforce the associations with cholesterol from the GLGC GWAS. This confirms the hypothesized phenotypic effect of the selected variants in terms of increased EPA and ARA levels of the putatively positively selected variants in the European population. We also tested whether the effect of dietary intake of EPA on cholesterol levels differed across genotypes of rs174594. Interestingly, we observed that the slope of the relationship between EPA intake and total cholesterol decreased with each copy of the derived allele (from 53.9 mg/dL per g/d of intake for CC homozygotes [95% CI: 15.2-92.7] to -3.49 for AA homozygotes [95% CI: -21.7-14.7], interaction p-value=0.02). Results for LDL cholesterol were similar (Table 4). This suggests individuals with the derived allele appear to be somewhat protected from the negative effects of increased EPA intake on plasma cholesterol levels.

Taken together, the NHS/HPFS and the ADDITION-PRO GWAS results provide evidence that alleles selected in Europe are associated with a decrease in linoleic acid levels and an increase in eicosapentaenoic acid, suggesting a more rapid conversion of SC-PUFAs to LC-PUFAs.

**Table 4.**
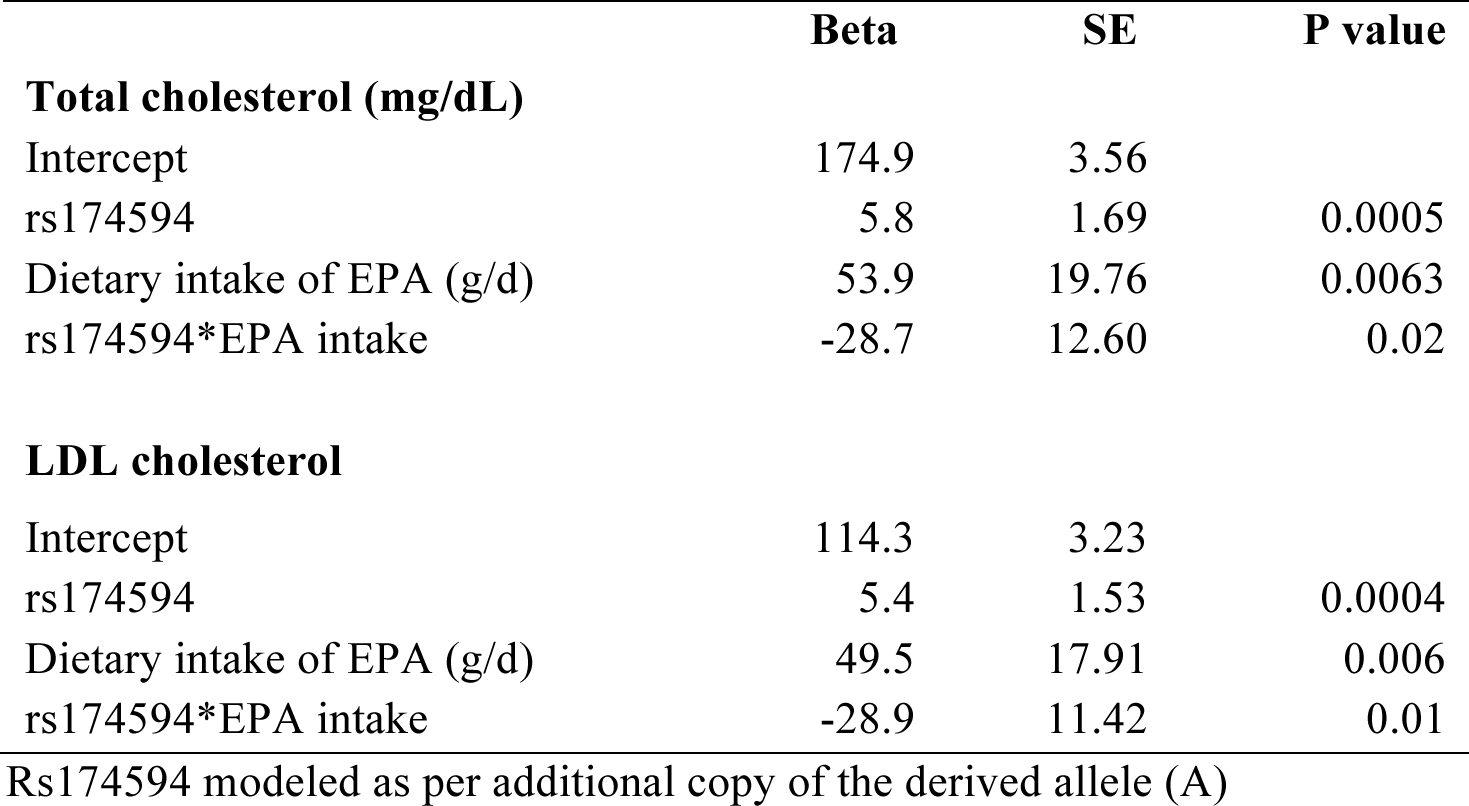
Interaction between dietary intake of EPA and rs174594 on plasma cholesterol and LDL cholesterol levels in the Nurses Health Study and Health Professionals Follow-Up Study.

|  | Beta | SE | P value |
| --- | --- | --- | --- |
| **Total cholesterol (mg/dL)** |  |  |  |
| Intercept | 174.9 | 3.56 |  |
| rs174594 | 5.8 | 1.69 | 0.0005 |
| Dietary intake of EPA (g/d) | 53.9 | 19.76 | 0.0063 |
| rs174594*EPA intake | -28.7 | 12.60 | 0.02 |
| **LDL cholesterol** |  |  |  |
| Intercept | 114.3 | 3.23 |  |
| rs174594 | 5.4 | 1.53 | 0.0004 |
| Dietary intake of EPA (g/d) | 49.5 | 17.91 | 0.006 |
| rs174594*EPA intake | -28.9 | 11.42 | 0.01 |

Rs174594 modeled as per additional copy of the derived allele (A)

## Discussion

Analyzing sequencing data from Bronze Age individuals allowed us to obtain a high-resolution picture of allele frequency changes in the *FADS* gene cluster. Mathieson, *et al*. (2015) previously showed that selection has acted in the *FADS* region in Europe since the Bronze Age. The lead SNP from that study (rs174546) is located in *FADS1* and could be a target of selection. However, the observed changes in allele frequency in Europe through time are best explained by selection acting on one of the HDSs in FADS2, or on a combination of these SNPs and the previously identified indel (rs66698963). Interestingly, rs174546 shows larger allele frequency differences between present-day Northern and Southern European than between present-day and Bronze Age Europeans, suggesting that selection has acted differentially in Northern and Southern Europe. In fact, it is possible that selection has acted on multiple SNPs, and in different regions of Europe in different ways. On a more global scale, we observe that selective pressures in Europe, Greenland, Africa, and South Asia have driven allele frequency changes in different SNPs that are only in weak LD with each other. We also find evidence of allelic heterogeneity in *FADS1* and *FADS2*. Presumably, there are many SNPs in the region affecting fatty acid desaturase activity that can serve as substrates for selection under varying environmental/dietary conditions. We also observe that the *FADS* region harbors highly differentiated haplotypes with genetic variability dating to much older times than the split from Denisovans and Neanderthals, possibly suggesting that long term balancing selection has been acting in this region.

The alleles with the largest allele frequency changes in Europe are strongly associated with expression changes in *FADS1* and *FADS2*, and are also associated with multiple lipid-related phenotypes: in particular, reduced linoleic acid levels. This is exactly the opposite pattern as observed for the selected alleles in the Inuit (Fumagalli, *et al*. (2015), a population where it is hypothesized that selection acted to decrease conversion of SC-PUFAs to LC-PUFAs to compensate for the relative high dietary intake of LC-PUFAs. Fumagalli, *et al*. (2015) showed that the favored allele in Inuit was associated with increased levels of linoleic acid and α-linolenic acid, and decreased levels of arachidonic acid and eicosapentaenoic acid. However, in Europe, selection has favored alleles associated with a decrease in linoleic acid levels and an increase in eicosapentaenoic acid, potentially due to an improved capacity to metabolize LC-PUFAs from SC-PUFAs. We hypothesize that this may be an adaptation to a diet rich in fatty acids derived from plant sources, but relatively poor in fatty acids derived from fish or mammals. The introduction and spread of agriculture in Europe likely produced a radical dietary shift in populations that embraced this practice. Agricultural diets would have led to a higher consumption of grains and other plant-derived foods, relative to hunter-gatherer populations. Alleles that increase the rate of conversion of SC-PUFAs to LC-PUFAs would therefore have been favored.

The fact that strong selection is acting on common alleles segregating in Europeans in the *FADS* region suggests that these alleles may be very important in determining the relative nutritional benefits of diets differing in their fatty acid composition. There has been substantial debate on the possible benefits of a high intake of PUFAs. It is possible that variants in the *FADS* region may underlie individual differences in optimal dietary fatty acid profiles. If so, the *FADS* region variants might help guide the development of individualized diets informed by genomics. However, functional studies of the region, possibly in cell cultures, are needed to determine the effects of individual SNPs on desaturase activity. Likewise, more epidemiological studies are needed to identify possible interactions between genetic variants in the *FADS* region and diet, and their joint effect on human health in general.

## Materials and Methods

### Haplotype network and structure analysis

First, we used Haploview v4.1 (Barrett *et al*. 2005) to produce linkage disequilibrium (LD) plots using the SNP data from the HapMap project (International HapMap 3 Consortium 2010) (Figure 3, Suppl. Figure 1). We selected the 30 most frequent haplotypes from the 1000 Genomes data (1000 Genomes Project Consortium 2015) in each of the two main LD blocks in the *FADS* region. We then plotted them in a haplotype network using the R package pegas (Paradis *et al*. 2010) along with the Altai Neanderthal (Prüfer *et al*. 2014) and Denisova (Meyer *et al*. 2012) haplotypes (Figure. 3, Suppl. Fig. 3). We also plotted the haplotype structure of the two blocks in each of the 1000 Genomes super-populations. We selected SNPs with more than 5% minor allele frequency in each super-population. We then ordered the resulting haplotypes by decreasing similarity to either the Denisova or the Altai Neanderthal genome, using Hamming distances, as in Racimo *et al*. (2015).

### Estimation of Allele Frequency Changes

Allele frequencies changes were calculated in R (version 3.1.2) by taking the absolute values of the allele frequency differences across the *FADS* region (chr11: 61567097-61659006) between two population panels. We used the 1000 Genomes Project Phase 3 data (Genomes Project, *et al*. 2015), specifically the five populations of European ancestry CEU, GBR, FIN, IBS, and TSI (often combined into a super population denoted as EUR), and an ancient DNA data set comprised of low-coverage (0.7x average) sequencing data from 101 Bronze Age Eurasians (Allentoft, *et al*. (2015). In several comparisons, we used a subset of the Bronze Age Eurasians dataset, consisting of 57 Bronze Age Europeans whose samples were collected west of the Ural Mountains.

### Expression data

All eQTL data was obtained through queries of the GTEx Release V6 (dbGaP Accession phs000424.v6.p1) (Lonsdale, *et al*. 2013).

### CAVIAR analyses

CAVIAR (CAusal Variants Identification in Associated Regions) is a probabilistic method that was originally developed to detect a confidence set of SNPs that contains all the causal variants with a predefined probability ρ (e.g., 90% or 95%) in a locus (Hormozdiari *et al*. 2014) taking local LD into account. Recently, we extended CAVIAR to detect allelic heterogeneity (Hormozdiari *et al*. forthcoming). Allelic heterogeneity (AH) is a phenomenon where more than one variant simultaneously affects a phenotype. CAVIAR incorporates the observed marginal statistics and LD structure to detect loci that harbor AH. Levering the fact that the marginal statistics follow a MVN distribution given the set of variants that are causal, we can quantify the probability of having a certain number of causal variants in a locus. We have shown that CAVIAR (Hormozdiari *et al*. in preparation) has a low false negative rate in detecting loci that harbor AH even when the true causal variants are uncollected. Thus, CAVIAR most likely accurately detects true loci that harbor AH.

### Association Analyses - ADDITION-PRO

The ADDITION-PRO study (2009-2011) (Johansen, et al. 2012) is a continuation of the Danish arm of the ADDITION study (2001-2006) (Lauritzen, et al. 2000), in which individuals with normal glucose tolerance (NGT), impaired fasting glycaemia (IFG), impaired glucose tolerance (IGT) or type 2 diabetes are followed. A total of 2082 participants have undergone an extensive examination, including detailed characterization of glycemic status based on a 3-point OGTT. Ethical approval was obtained from the Scientific Ethics Committee of the Central Denmark Region (journal no. 20080229) and all participants provided oral and written informed consent before participating in the study. Genotyping was performed using the Illumina Infinium HumanCoreExome Beadchip platform (Illumina, San Diego, CA) and genotypes were called using the Genotyping module (version 1.9.4) of GenomeStudio software (version 2011.1, Illumina). Blood samples for the measurement of lipid-related phenotypes were collected in EDTA tubes and put on ice immediately, centrifuged to collect the plasma content, and stored at -80 degree. Samples were taken out from -80 degree and thawed overnight in a refrigerator prior to sample preparation. Proton nuclear magnetic resonance (NMR) spectroscopy was used for measurement. NMR methods have been described in detail previously (Kettunen, et al. 2016). Genotype-phenotype correlations were studied with PLINK version 1.07 (Purcell, et al. 2007) using linear regression analyses adjusted for age, sex, BMI and glucose level at the time point.

### Association Analyses - NHS/HPFS

The Nurses’ Health Study (NHS) is an ongoing prospective cohort study of 121 700 female registered nurses from 11 US states who were 30 to 55 years of age at study initiation in 1976. The Health Professionals Follow-Up Study (HPFS) is a similar prospective study consisting of 51 529 male health professionals from all US states who were 40 to 75 years of age at baseline in 1986. For both cohorts, mailed questionnaires were administered biennially to assess lifestyle factors and health status, with a follow-up rate exceeding 90% for each 2-year cycle. Between 1989 and 1990, a total of 32 826 NHS participants provided a blood sample. With a similar protocol, blood samples were collected from 18 225 HPFS participants between 1993 and 1995. In both cohorts, the blood samples were returned via an overnight courier, and most (≥95%) of the blood samples arrived within 24 hours of phlebotomy. On arrival, samples were centrifuged and divided into aliquots in cryotubes as plasma, buffy coat, and erythrocyte fractions, which were then stored in liquid nitrogen freezers at −130°C or colder until analysis. We analyzed 2,288 European-ancestry women and men who were free of CHD at time of blood collection in the NHS and HPFS. These subjects were selected as a nested case-control study of CHD (n=342 cases for NHS and n=435 cases for HPFS) and were genotyped using the Affymetrix 6.0 SNP array (Jensen, et al. 2011). Genotypes were imputed using MACH and the 1,000 Genomes Project Phase 1 v3 reference panel. Plasma fatty acid concentrations were analyzed by gas-liquid chromatography as described previously by Malik, et al. (2015). The concentration of each fatty acid was expressed as a percentage of the total fatty acid content. Concentrations of total cholesterol and high-density lipoprotein cholesterol (HDL-C) were measured with a Hitachi 911 analyzer using reagents and calibrators from Roche Diagnostics (Indianapolis, IN). Low-density lipoprotein cholesterol levels were determined with a homogenous direct method from Genzyme (Cambridge, MA).

Association analyses were conducted using linear regression, regressing fatty acid and lipid measurements on genotype dosages. For interaction analyses, we estimated dietary intake of EPA with validated food frequency questionnaires inquiring about intake over the year prior to blood draw (1990 in NHS and 1994 in HPFS).

## Acknowledgments

The Nurses’ Health Study (NHS) and Health Professionals Follow-Up Study (HPFS) were supported by UM1 CA186107, HL34594, CA87969, CA49449, HL35464, CA55075, R01 HL088521, CA186107, CA87969, CA49449, HL60712, and CA167552 from the National Institutes of Health, Bethesda, MD, with additional support for genotyping from Merck Research Laboratories, North Wales, PA.

